# LinkDTI: Drug-Target Interactions prediction through a Link Prediction framework on Biomedical Knowledge Graph

**DOI:** 10.64898/2026.02.21.707210

**Authors:** Madhurima Mondal, Subrahmanyam Arunachalam, Shili Wu, Aniruddha Datta

**Author notes:** Corresponding author: Madhurima Mondal.

## Abstract

Computational drug-target interactions (DTI) prediction serves as a valuable tool for drug discovery and repurposing by cost-effectively narrowing down the potential drug-target space. This paper presents LinkDTI, a computational framework that predicts DTIs by identifying connections within a heterogeneous knowledge graph of drugs, proteins, diseases, and side effects. Unlike methods that rely on mathematical techniques like matrix completion or similarity-based scoring, LinkDTI uses an advanced graph-based approach to capture relationships between biomedical entities. Specifically, LinkDTI applies a modified version of the multilayer GraphSAGE model that learns from the heterogeneous knowledge graph and predicts potential drug-target interactions. Our model incorporates negative sampling that balances the data to address the issue of having more negative than positive interactions. Our results show that LinkDTI consistently performs better in AUROC and AUPRC than baseline methods by at least 2.5% across different sampling ratios and conditions. Subsequently, it identifies approximately 945 new potential DTIs, marking a 49.14% increase over known DTIs. Overall, LinkDTI offers a simple yet effective method for integrating diverse biomedical data to identify potential drug-target interactions. The code and data can be found at https://github.com/hub2nature/LinkDTI_heterogenous_KG.git.

## I. INTRODUCTION

Drugs interact with target proteins to produce effective therapeutic outcomes [1]. Therefore, predicting drug-target interactions (DTIs) is pivotal in drug discovery, development, and repurposing [2]. Given millions of potential chemical compounds, it is impractical and infeasible to conduct laboratory experiments to predict the DTIs for all of them [3]– [5]. Thus, despite recent developments in medical science, predicting DTI remains challenging. As a result, researchers have developed computational strategies that narrow down the possible drug-target space to predict DTIs [3], [6], [7]. Among these strategies, network-based approaches are particularly significant for their capacity to integrate diverse biomedical information [8]–[10]. In this paper, we aim to improve the accuracy of DTI prediction with a novel graph-based method that integrates relationships between drugs, target proteins, associated diseases, and corresponding side effects to capture the complex interactions that underlie drug-target interaction dynamics.

We propose LinkDTI, a multilayer heterogeneous GraphSAGE-based model followed by a link, i.e., edge prediction module, to effectively integrate the mentioned complex relationships [11], [12]. We utilize interaction matrices containing drugs, proteins, diseases, and side effects to create a heterogeneous knowledge graph. Unlike homogeneous graphs, which treat all nodes and edges uniformly, our model preserves distinctions between drugs, proteins, diseases, and side effects. The multilayer GraphSAGE mechanism allows the model to integrate node information from neighbors at multi-hop path lengths [13], [14]. Our link prediction strategy helps directly learn the node-node interaction probability, moving beyond traditional matrix completion or similarity-based scoring methods. Furthermore, we integrate a negative sampling strategy to balance highly skewed data and improve the prediction accuracy [8], [9], [15], [16]. This study provides a simpler and more effective framework for inferring meaningful interactions from diverse biomedical data.

Early machine learning models lay the groundwork for computational DTI predictions [6], [7]. These models cover diverse strategies such as similarity or distance scores, clusters, and matrix factorization. Similarity-based methods such as MSCMF often calculate drug-drug, target-target, and drug-target similarity scores from their chemical, functional, or sequential properties [17]. Methods like SITAR and SRP utilize a logistic regression to predict DTIs from various drug-drug and target-target similarities [18], [19]. NetLapRLS presents a semi-supervised learning method to address the challenge of handling the scarcity of known interactions and the vast number of unknown ones [16]. Distance-based methods like Nearest Neighbor (NN) use metrics such as Euclidean distance to measure the similarity between drugs and targets for DTI prediction [20]–[22]. Another technique, WNN-GIP (Weighted Nearest Neighbors with Gaussian Interaction Profile), combines nearest neighbor algorithms with Gaussian Interaction Profiles to enhance interaction predictions [23]. Probabilistic Matrix Factorization (PMF) decomposes the DTI matrix using collaborative filtering [24].

Learned feature-based methods first convert each drug-protein pair into fixed length vector, and use supervised learning to predict unknown DTIs. Support vector machines (SVMs) and ridge regression predict DTI labels by constructing hyperplanes from extracted features [25]– [27]. Ensemble methods like Random Forests and Bagging-based Ensemble (BE-DTI) utilize multiple classifiers to enhance DTI predictions [28], [29]. Furthermore, models like DrugRPE, integrate random projection and regression trees, while PUDT addresses positive-unlabeled learning scenarios to predict interactions effectively using weighted SVMs [30], [31]. Deep learning techniques such as DeepDTI first convert raw drug and protein inputs into embedded vectors and then apply deep belief networks to predict DTIs [32]. DeepConv-DTI predicts DTIs using convolution neural networks to capture drug and protein feature patterns [33]. DeepDTA leverages character-level representations of drugs and targets, while the more advanced method, MolTrans, utilizes transformer architecture and a convolution network to capture pair-wise drug-target features [34], [35]. DTI-CNN employs a Jaccard similarity coefficient and a random walk with restart to extract features from heterogeneous networks. This method can predict DTIs accurately using a convolution neural network but faces potential information loss during dimensionality reduction [36]. Recently developed methods such as DrugRepoAlign, Lint, and Y-Mol have contributed to DTI prediction using the latest Large Language Models (LLMs) by incorporating contextual embeddings to capture intricate relationships in electronic clinical records and medical articles [37]–[39].

Recent advances in graph embedding methods have demonstrated groundbreaking performances in graph data analysis [6]. A DTI prediction task can be compared to a link or edge prediction in a graph, where drugs and proteins are nodes, and their interaction labels are typically edges. Drug-target interaction matrices are generally used to construct such graphs. Node features are converted into low-dimensional vectors that can be utilized for DTI prediction. A heterogeneous graph-based method captures biological relationships by constructing a heterogeneous network from diverse biological data such as drug, protein, and chemical properties. One popular graph-based method, NeoDTI, aggregates neighborhood information followed by reconstructing the original network [9]. Although NeoDTI can capture accurate information, it solely dependents on matrix manipulation techniques that prevent NeoDTI from accurately collecting higher-order information [9]. DTINet converts a heterogeneous network into a low-dimensional homogeneous feature space which may ignore the vital heterogeneity of nodes that contribute to DTI predictions [10]. Like DTINet, GADTI also converts the heterogeneous network into a homogeneous space, followed by a random walk with restart to gather higher order information and matrix reconstruction [8]. These may limit GADTI with the loss of heterogeneous context and increased computational complexity. TriModel performs tensor decomposition to predict DTI from a knowledge graph. However, this method faces challenges with handling noisy data [40]. HampDTI integrates diverse biological data by utilizing a probabilistic matrix factorization. This model simplifies drug-target interaction prediction but lacks type-specific information in the network [41].

Thus, matrix manipulation or homogeneous representation-based methods for DTI prediction suffer from limitations such as high computational complexity, and difficulty capturing heterogeneity. LinkDTI overcomes these limitations by leveraging a Heterogeneous GraphSAGE network with multilayer message passing and edge prediction mechanisms. LinkDTI ensures a more promising understanding of complex biomedical relationships through type-specific distinctions among drugs, proteins, diseases, and side effects. We emphasize distinct connectivity among heterogeneous edges. Additionally, LinkDTI’s edge prediction strategy, combined with a robust negative sampling approach, provides a more scalable and adaptable framework for DTI prediction. The paper is organized as follows. In Section II formulates the drug-target interaction (DTI) prediction task as a link prediction problem in knowledge graphs. Section III describes knowledge graph construction, multilayer graph message passing, and the link prediction mechanism. We present datasets and the experimental setup in Section IV. The results obtained, and the performance comparison against existing state-of-the-art methods are presented in Section V. Section VI concludes the paper and suggests potential future work.

## II. DTI AS A LINK PREDICTION PROBLEM IN A KNOWLEDGE GRAPH

### KNOWLEDGE GRAPH

DTI prediction can be formulated as a link prediction task in a knowledge graph. This formulation aims to predict missing links (edges) between nodes representing biomedical entities such as drugs, proteins, diseases, and side effects. This representation leverages the structural relationships and node features to uncover potential interactions.

The heterogeneous knowledge graph is defined as *G* = (*V, E, R*), where:

- *V* = {*v*_1_, *v*_2_, …, *v*_*n*_}, with *v* ∈ {drug, protein, disease, side effect}, representing the set of nodes.
- *E* = {(*u, v, r*) | *u, v* ∈ *V, r* ∈ *R*}, the edges representing observed relationships between nodes.
- *R*, the set of binary-labeled relations, describing the semantics of edges, such as “drug-drug interaction” or “drug-disease association”.

Each edge (*u, v, r*) ∈ *E* indicates a relationship of type *r* ∈ *R* between the nodes *u* and *v*. Here, *r* ∈ {0, 1} where *r* = 1 denotes the presence of a relation, and *r* = 0 denotes the absence of a relation between the nodes. This binary representation aids in capturing both potential (missing) and existing linkages.

#### A. NODE REPRESENTATION

Each node *v* ∈ *V* is associated with a feature vector *h*_*v*_ ∈ ℝ^*d*^, where *d* is the dimensionality of the feature space. The feature matrix for all nodes is represented as:

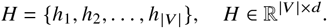

#### B. LINK PREDICTION

The link prediction task involves predicting whether a link (*u, v, r*) exists in the KG, i.e., determining whether:

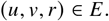

This can be formulated as learning a scoring function:

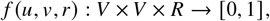

where *f* (*u, v, r*) outputs the likelihood of the edge (*u, v, r*) existing in *G*. The objective is to predict:

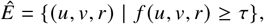

where *τ* is a threshold defining the decision boundary for link prediction.

The output function can be expressed probabilistically as:

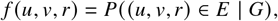

where *P* ((*u, v, r*) ∈ *E* | *G*) represents the probability of the edge’s existence in *G*. The goal of *f u, v, r* is to approximate the true likelihood of the existence of an edge, which relates to the actual knowledge graph structure.

## III. METHODS

### A. GRAPH CONSTRUCTION

We construct a heterogeneous knowledge graph *G* = (*V, E, R*) representing diverse biomedical relationships. The node set, *V* consists of drugs, proteins, diseases, and side effects. The set of edges, *E* contains edges: drug-drug interaction, protein-protein interaction, drug-protein interaction, drug-side effect association, drug-disease association, and protein-disease association. Drug-drug interaction, proteinprotein interaction, and drug-protein interaction are marked as undirected links, whereas other edges are marked as directed links. Labels in *R* represent whether two nodes are connected or not to capture node relationships.

### B. NODE INITIALIZATION

Each node in *G* is initialized with random feature vectors of dimension *d* sampled from a normal distribution :

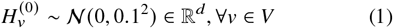

### C. HETEROGENEOUS GRAPHSAGE ARCHITECTURE

Our knowledge graph stores heterogeneous node information. We utilize a three-layer GraphSAGE architecture to update the node features with meaningful information. We modify the conventional GraphSAGE by removing the self-loops of each node while updating the node features. We add this modification to ensure that updated features are influenced solely by the information from neighboring nodes. This forces the model to rely on the structural context provided by neighboring nodes and avoid overfitting on its raw node features. Each layer of the GraphSAGE network employs a mean-based aggregation mechanism for propagating and collecting neighborhood information. The aforementioned knowledge graph contains multiple edge types that represent interactions or associations between nodes, such as drug-drug interaction, drug-disease association. We apply a modified version of GraphSAGE for each edge type, where the first GraphSAGE layer computes the immediate node embeddings as follows:

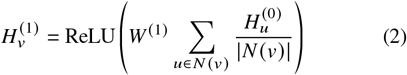

where *N* (*v*) represents the neighbors of *v* and *W* ^(1)^ is the learnable weight matrix for the first layer. The model repeats this process for subsequent layers to capture higher-order information. The final layer produces the node embeddings 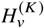 through *K* layers such that,

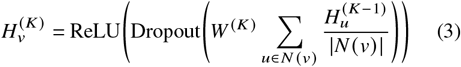

### D. EDGE REPRESENTATION AND LINK PREDICTION

To predict an edge (*u, v, r*) between source *u* and target *v*, we concatenated their embedded features

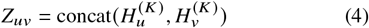

where *K* is the total number of layers considered in the model. An edge prediction function *f* is applied using a Multilayer Perceptron (MLP):

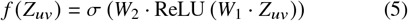

where *Z*_*uv*_ is the concatenated feature vector of nodes *u* and *v, W*_1_ and *W*_2_ are trainable weight matrices, ReLU (·) is the Rectified Linear Unit activation function, σ(·) is the sigmoid function, ensuring the output is a probability. The proposed architecture is represented in Fig 1.

**FIGURE 1:**
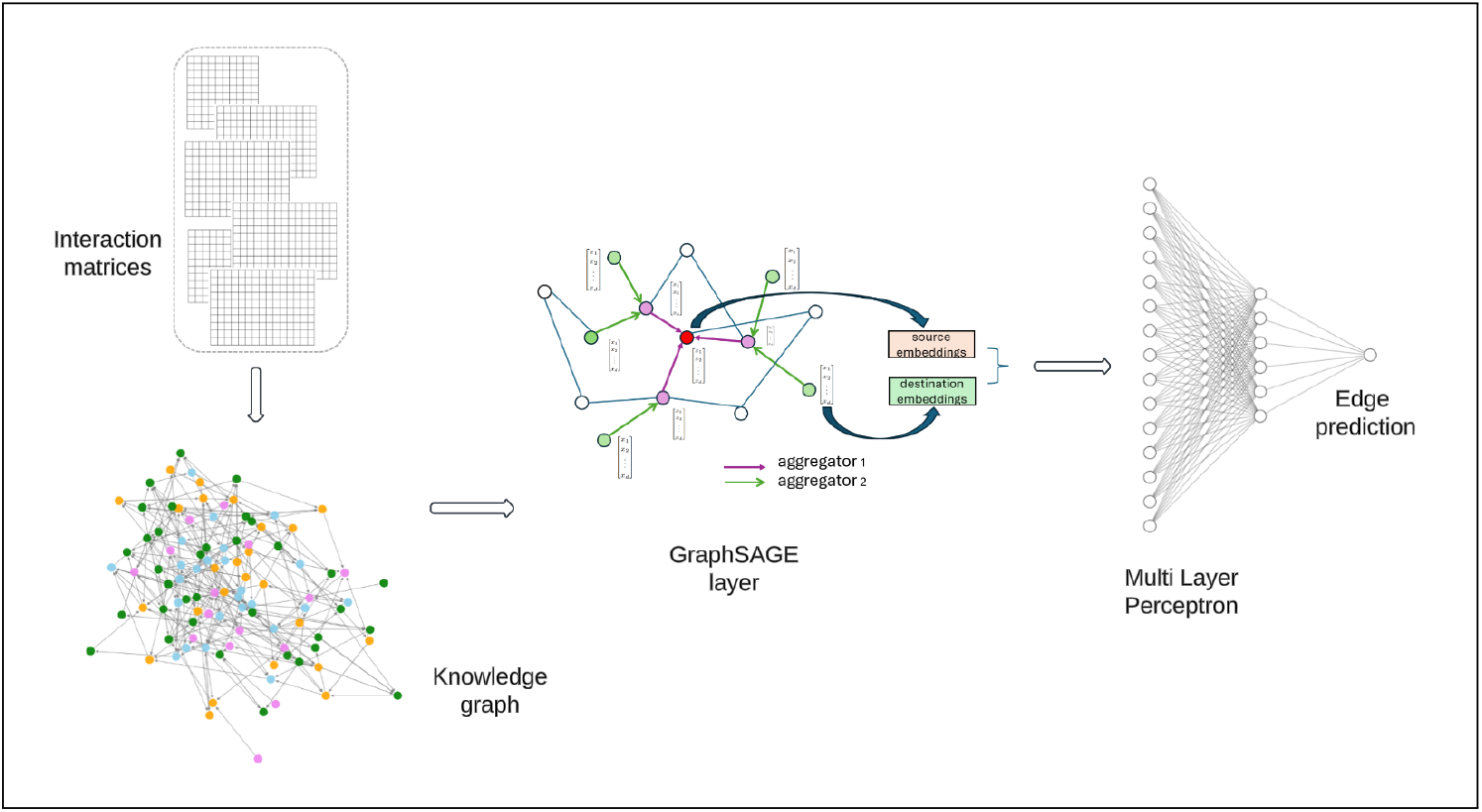
Block diagram of the LinkDTI. In the figure interaction matrices include drug-drug, drug-protein, protein-protein, drug-disease, drug-side effect, and protein-disease associations, which are used for knowledge graph construction. The multilayer Heterogeneous GraphSAGE architecture captures neighborhood information, and the edge representation module concatenates node embeddings for link prediction using a multilayer perceptron (MLP).

## IV. MATERIALS

### A. DATASET

In this paper utilizes publicly available datasets from previous studies [8]–[10]. The datasets contain four node types, i.e., drugs, targets, diseases, and side effects, and six edge types between the different nodes, such as drug-drug interaction, protein-protein interaction, drug-protein interaction, protein-drug interaction, drug-side effect association, side effect-drug association, drug-disease association, disease-drug association, protein-disease association, disease-protein association. We summarize the node and edge statistics in Table 1 and Table 2, respectively. The available data are curated from well-established public databases. For example, drug–target, and drug-drug interactions are sourced from DrugBank v3.0 [42]. Protein-protein interactions are obtained from the Human Protein Reference Database (HPRD) Release 9 [43]. Drug–disease, and protein–disease association data are extracted from the Comparative Toxicogenomics Database [44]. Information on drug side effects is derived from SIDER Version 2 [45]. The drug–side effect, and side effect–drug, associations differ in direction and query context. Modeling both directions in the knowledge graph enhances its representational power.

**TABLE 1:** Node Statistics.

| Node Type | Count |
| --- | --- |
| Drug | 708 |
| Protein | 1,512 |
| Disease | 5,603 |
| Side Effect | 4,192 |
| Total | 12,015 |

**TABLE 2:** Edge Statistics.

| Edge Type | Count |
| --- | --- |
| Drug-Target Interaction | 1,923 |
| Drug-Drug Interaction | 10,036 |
| Protein-Protein Interaction | 7,363 |
| Drug-Disease Association | 199,214 |
| Drug-Side Effect Association | 80,164 |
| Protein-Disease Association | 1,596,745 |
| Total | 1,895,445 |

### B. EXPERIMENTAL SETUP

We construct the knowledge graph *G* representing diverse relationships by preprocessing the interaction matrices to evaluate the proposed framework. We represent symmetric interactions like drug-drug and protein-protein as undirected, asymmetric associations such as drug-disease and drug-side effect as directed to preserve the contextual directionality of these relationships. We initialize each node with random feature vectors sampled from a normal distribution mentioned in Eq. 1, with dimension *d* = 512. The model generates negative samples from unconnected pairs in the drug-target interaction matrix. To ensure a balanced dataset, we randomly select a subset of these negatively sampled pairs proportional to the number of positive samples. We maintain distinct canonical edge types and generate interaction-specific samples and labels for each interaction type (e.g., drug-drug, protein-disease). We preserve this heterogeneity to help the model capture the unique relationships between biomedical entities while maintaining their structural relationship within *G*. For training; we create training, validation, and test subgraphs from *G* by dividing the input dataset into 80% training data, 10% validation data, and 10% test data. We employ stratified splitting to ensure balanced and disjoint splits across edge types in the dataset, leading to disjoint subgraphs. Node embeddings are updated using a three-layer heterogeneous GraphSAGE model (Eq. 3) followed by a dropout of 0.5 to avoid overfitting and a ReLU layer as an activation function. We pass the final concatenated source and target embeddings of an edge through a three-layer Multilayer Perceptron (MLP) with hidden dimensions [256, 128, 1]. To receive edge probabilities, we apply a sigmoid function at the output. A brief description of the experimental set is portrayed in a flow chart in Figure 2.

**FIGURE 2:**
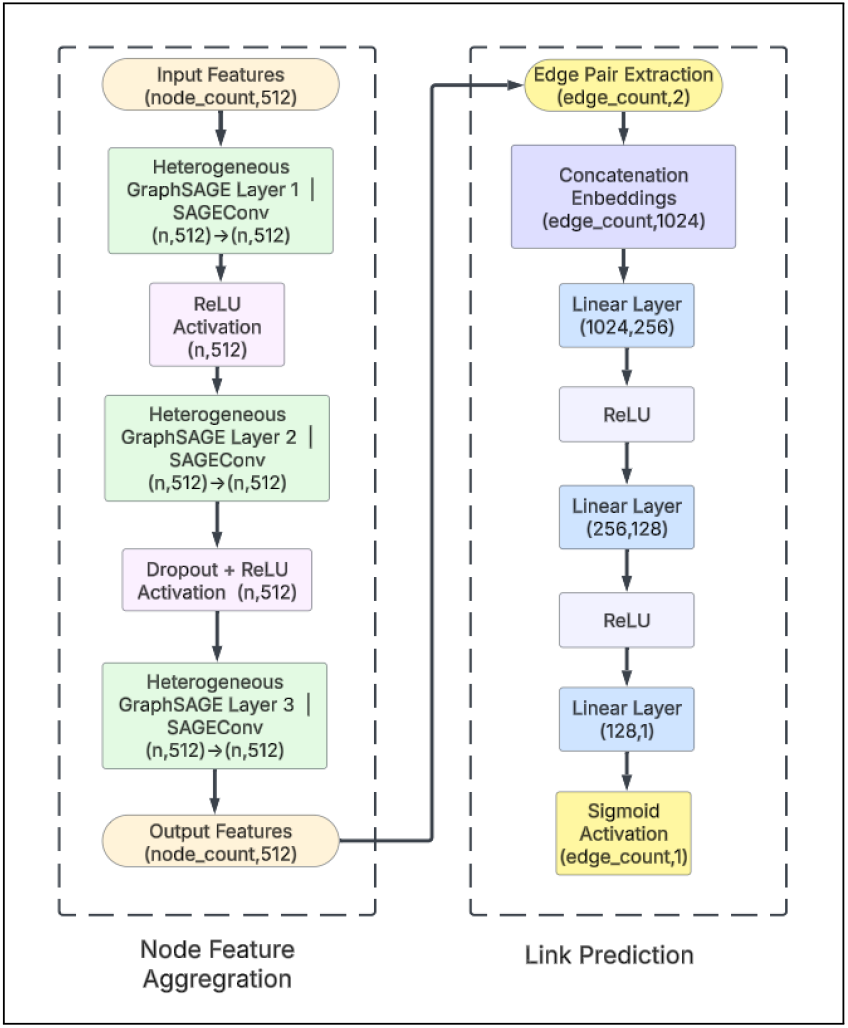
A visual interpretation of the LinkDTI process through a flow chart.

We initialize the weights of all layers using Xavier uniform initialization before training [46]. This enables the neural network layers to start the parameters with well-suited values and stabilize the training process. The model employs the Adam optimizer with an initial learning rate of 0.001. Binary Cross-Entropy (BCE) is the loss function to minimize prediction error. Training consists of edge processing in batches of size 512, mixed by positive and negative samples for the given edge type. Batch training ensures efficient computation while effectively leveraging GPU memory. We apply early stopping with a patience of 9 epochs. The paper evaluates the model using the Area Under the Receiver Operating Characteristic Curve (AUROC) and the Area Under the Precision-Recall Curve (AUPRC) metrics [47], [48]. We averaged performance over five random runs to ensure robustness and reproducibility. All experiments were performed on an NVIDIA A100 GPU with 40 GB of memory.

### C. EVALUATION METRICS

We utilize two widely used evaluation metrics, the Area Under the Receiver Operating Characteristic Curve (AUROC) and the Area Under the Precision-Recall Curve (AUPR), for the classification task in our paper to assess our model’s performance [47], [48]. AUROC measures a model’s ability to discriminate between positive and negative samples by plotting the true positive rate (TPR) against the false positive rate (FPR) [47]. AUPRC indicates the trade-off between precision and recall [49]. As it assesses finding genuine positive interactions and reducing false positives, a higher AUROC value indicates accurate classification. AUROC and AUPRC are useful in this study for dealing with a classification task within an imbalanced dataset.

## V. RESULTS

To evaluate the performance of our proposed model, we performed various experiments and compared the results with several baseline models. We provide experiments across varying dataset configurations such as balanced sample settings, 1:10 positive-to-negative sample ratios, treating all unknown samples as negative samples, and eliminating similar drugs and proteins [15]. Our model outperforms NetLapRLS [16], DTINet [10], DTI-CNN [36], NeoDTI [9], and GADTI [8] in AUROC and AUPRC values measured over five different runs. Our findings are given below:

### A. PERFORMANCE EVALUATION UNDER A 1:10 POSITIVE-TO-NEGATIVE SAMPLE RATIO

As described in Section IV-B, a setback to deal with drug-target interaction prediction is to handle highly imbalanced datasets with significantly less known interactions than unknown interactions. We first assess our model in a 1:10 positive-to-negative sampling ratio to factor in the class imbalance. The mean AUROC and mean AUPRC over five runs and the uncertainty are given in Figure 3. Compared to all the evaluation models, LinkDTI outperforms, achieving an AUROC of **0.9784** and an AUPRC of **0.9264**. LinkDTI surpasses the second-best model, GADTI, that achieved an AUROC of 0.9549 and an AUPRC of 0.8557, by 2.5% in AUROC and 7.6% in AUPRC. NeoDTI, with an AUROC of 0.9507 and an AUPRC of 0.8684, ranks third with a 2.8% lower AUROC and a 6.3% drop in AUPRC compared to LinkDTI. These findings highlight that, although GADTI and NeoDTI can effectively predict drug-target interaction, LinkDTI will be more reliable in DTI predictions. Despite leveraging convolutional architectures, DTI-CNN that achieves an AUROC of 0.9396 and an AUPRC of 0.8000, failed to match LinkDTI’s effectiveness. LinkDTI performes significantly better than traditional machine learning models such as NetLapRLS (AUROC = 0.9025, AUPRC = 0.7970) and DTINet (AUROC = 0.9140, AUPRC = 0.7630), which struggle with maintaining high precision in imbalanced data. The experiment with 1:10 positive-to-negative sampling ratio demonstrates LinkDTI’s superiority in differentiating interacting drug-target pairs from non-interacting pairs. LinkDTI is proven to be less prone to false negatives by achieving a high AUPRC value, which is a useful criterion in experimental drug discovery and drug repurposing.

**FIGURE 3:**
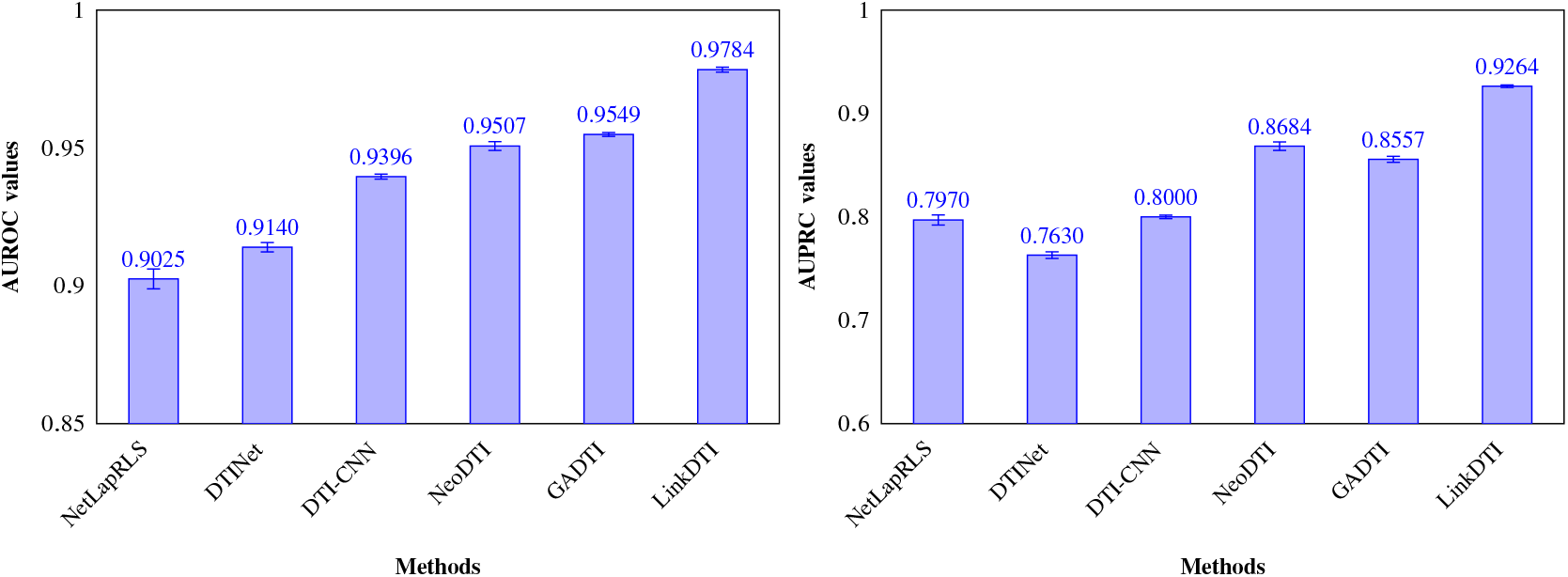
Comparison of AUROC and AUPRC values for different models in 1:10 positive to negative samples ratio.

### B. PERFORMANCE EVALUATION ON THE FULL DATASET WITH ALL UNKNOWN SAMPLES CONSIDERED

We analyze LinkDTI’s robustness by evaluating the model while considering all unknown DTIs in the dataset as negative samples. This setup, where the positive-to-negative sample ratio is roughly 555, is particularly valuable as it mimics the real-world scenario. Despite the increased challenge, LinkDTI performs quite well in a highly skewed dataset by achieving an AUROC of **0.9562** and an AUPRC of **0.8376**. As summarized in Figure 4, this experimental setup reinforces LinkDTI’s strength in achieving high recall and precision among all models. LinkDTI demonstrates superior performance with 2.3% improved AUROC and an AUPRC increase of 35.1%, than GADTI, that achieved an AUROC of 0.9345 and AUPRC of 0.6196. Similarly, compared to NeoDTI that results in an AUROC of 0.9123, and an AUPRC of 0.6012, LinkDTI outperformed by 4.8% in AUROC and 39.4% in AUPRC. These results indicate that GADTI and NeoDTI although maintained reasonable AUROC scores, they were prone to predict false positives, but LinkDTI could sustain the imbalanced dataset. Surprisingly, DTI-CNN showed a significant drop in performance in both AUROC (0.8764) and AUPRC (0.2963). This suggests that, when negative samples vastly outnumber its counterpart, convolution-based feature extraction may not be quite successful. NetLapRLS with AUROC of 0.9041 and AUPRC of 0.2837, and DTINet with AUROC of 0.9125 and AUPRC of 0.2800 performed poorly, demonstrating inefficacy in dealing with extreme class imbalance. This suggests that LinkDTI generalizes well with and without controlled training conditions.

**FIGURE 4:**
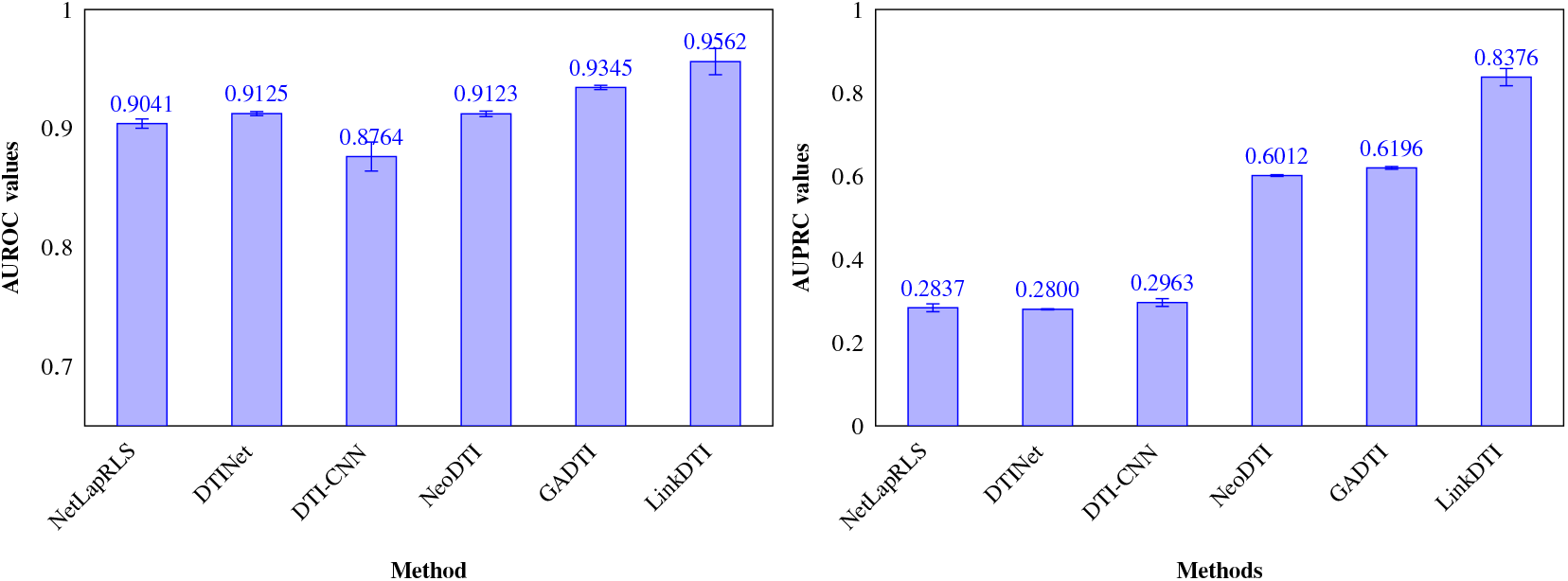
Comparison of AUROC and AUPRC values for different models when all the negative samples were considered in the study.

### C. PERFORMANCE ON A BALANCED 1:1 SAMPLE RATIO

To further investigate the effects of class imbalance on our model’s performance, we evaluate the LinkDTI under a balanced dataset configuration with a 1:1 positive-to-negative sampling ratio. This experimental setup significantly improves the AUPRC scores across all models as compared to the 1:10 sampling scenario. In this setup too, LinkDTI achieves the best results among other models with AUROC of **0.9850** and AUPRC of **0.9850**. LinkDTI defeats NeoDTI by approximately 4.23% in AUROC and 3.79% in AUPRC. Similarly, compared to GADTI, LinkDTI shows an improvement of around 4.89% in AUROC and 3.99% in AUPRC. DTINet, DTICNN, and NetLapRLS performed better than in the imbalanced setting in V-B, but still lags behind LinkDTI, reinforcing the idea that their feature extraction capabilities are not as sophisticated as graph-based methods like LinkDTI. The comparison plot is given in Figure 5.

**FIGURE 5:**
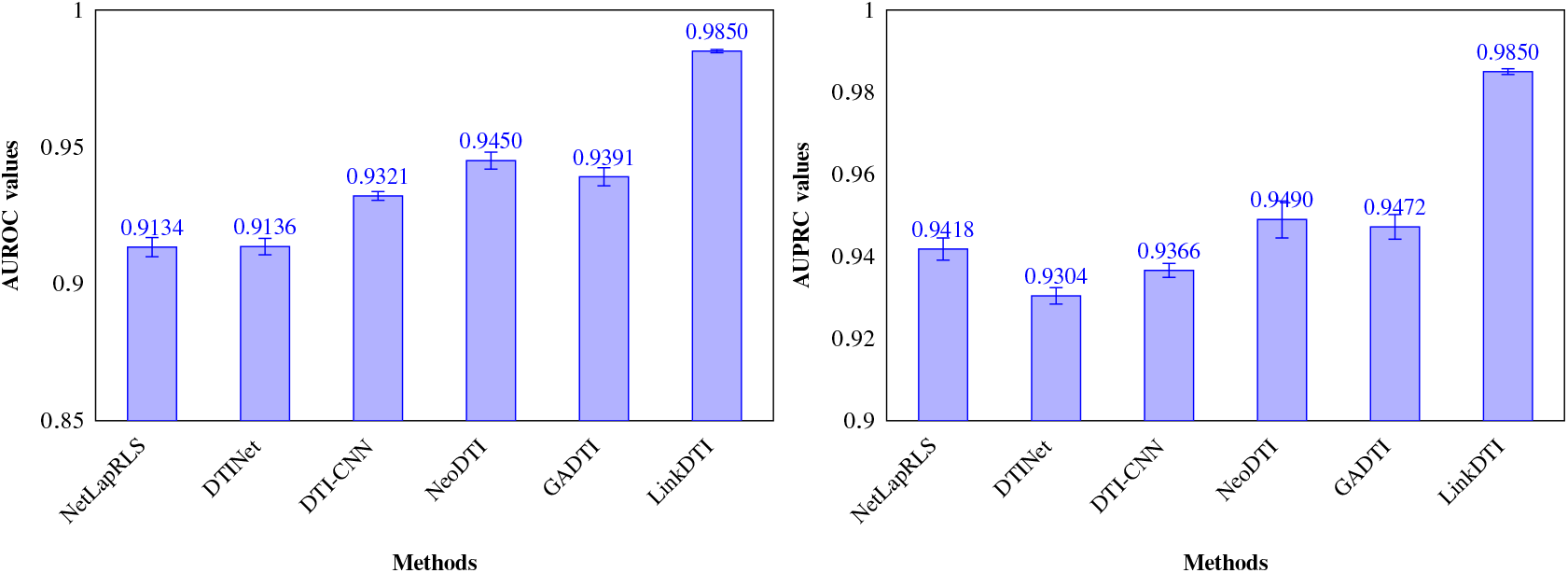
Comparison of AUROC and AUPRC values for different models in 1:1 positive to negative samples ratio.

### D. PERFORMANCE ON DATASET AFTER REMOVING DTIS OF SIMILAR DRUGS AND PROTEINS PAIRS

In addition to varying sample distributions, we examine LinkDTI’s performance when similar drugs and proteins were removed from the input drug-target interaction matrix. The reduced dataset, which includes 968 positive and 1,069,528 negative samples, contains fewer positive samples than the original DTIs. We considered a 1:10 positive-to-negative sampling on the dataset to wipe out the class imbalance to compare all models. LinkDTI achieves an AUROC of **0.9765** and an AUPRC of **0.9227** and outperforms DTI-CNN, which shows relatively strong performance, achieving an AUROC of 0.9333 and an AUPRC of 0.7936. Interestingly, GADTI and NeoDTI faced performance degradation in this reduced DTI setting compared to Section V-A. GADTI scored an AUROC of 0.9131 and an AUPRC of 0.7246, while NeoDTI records an AUROC of 0.8978 and an AUPRC of 0.6933. Unfortunately, NetLapRLS and DTINet exhibits the most significant performance drops. These patterns suggest that these models mainly depend on data diversity to properly differentiate between positive and negative interactions. The reduced DTI experiment indicates that the accuracy of LinkDTI sustains stability even in diverse setups. Figure 6 showcases a complete performance comparison of the models.

**FIGURE 6:**
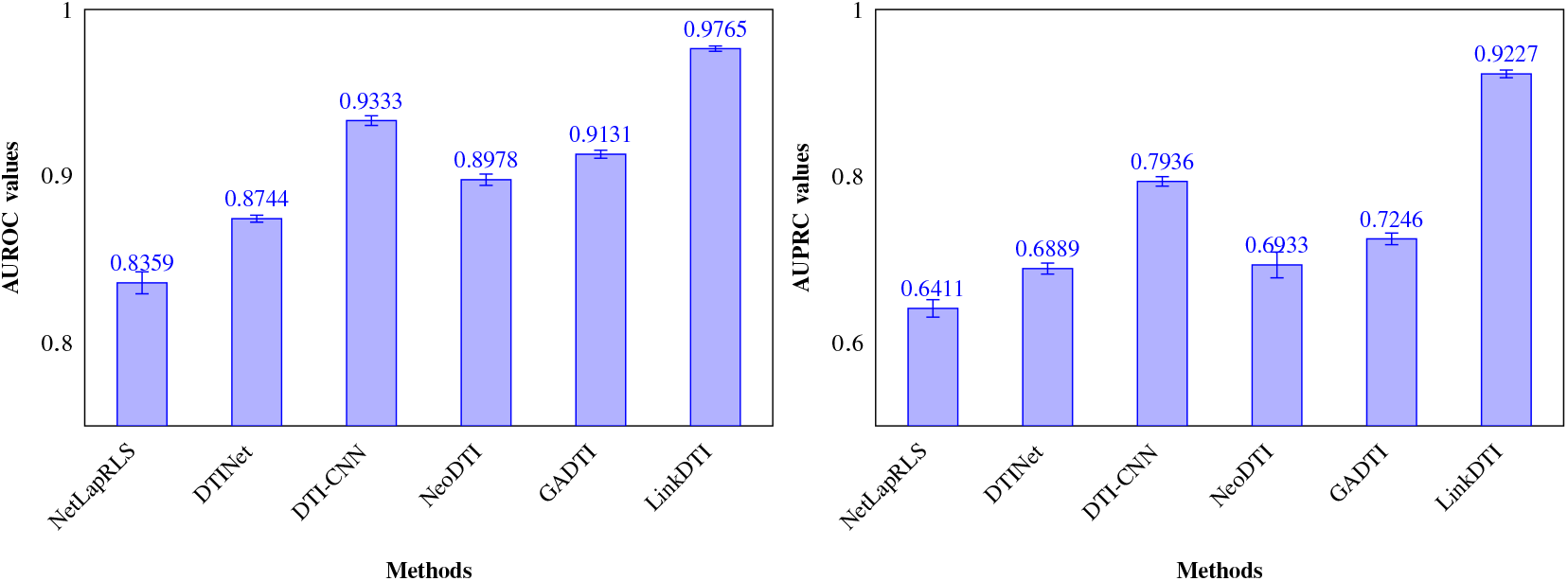
Comparison of AUROC and AUPRC values for different models in 1:10 positive to negative samples ratio and similar drugs and proteins removed.

### E. CASE STUDY ON NEWLY PREDICTED DTIS

To further evaluate LinkDTI’s performance, we extract the drug and protein embeddings from the trained and updated knowledge graph to generate a drug-target interaction matrix from the new node features. We binarize the DTIs using a range of threshold values. The cells in the newly created interaction matrix were categorized as 1 if their value is above the threshold and as 0 if their value is below it. To identify the newly predicted DTIs from the old ones, we remove all the known positive DTIs using the original drug-target interaction matrix as the input. We find the original drug-target interaction matrix had 1923 positive interactions. LinkDTI predicts around **945** new positive drug-target interactions, that is **49.14%** higher with a threshold of 0.745. The newly predicted DTIs suggest that LinkDTI is a potential and reliable tool for drug-target interaction prediction. Newly predicted DTIs over a range of threshold is given in Figure 7.

**FIGURE 7:**
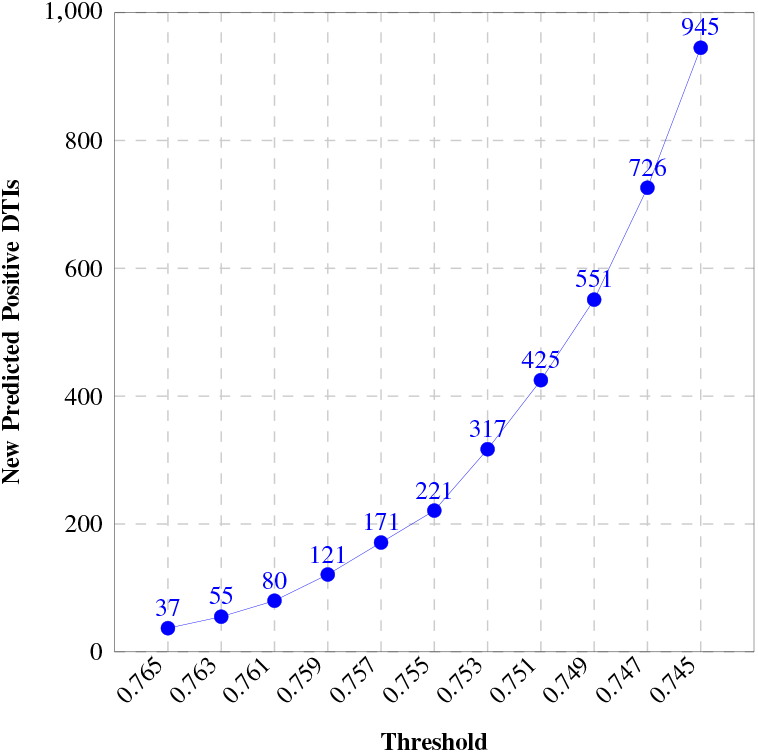
Newly predicted positive DTIs for different threshold values

## VI. CONCLUSION

In this study, we propose a robust framework for drug-target interaction (DTI) prediction. LinkDTI, leveraging a heterogeneous GraphSAGE network, integrates link prediction within the graph network. LinkDTI outperforms traditional machine learning models with matrix reconstruction methods such as as NetLapRLS, DTINet, DTI-CNNeoDTI,DTI, as well as homogeneous graph convolution-based approaches like GADTI by effectively capturing the heterogeneity among biomedical entities like drugs, target proteins, diseases, and side effects. This suggests that the inclusion of heterogeneity in the graphSAGE has demonstrated strong performance improvement. AUROC and AUPRC achieved by LinkDTI in multiple experimental setups suggested its robustness and accuracy. LinkDTI’s performance in maintaining high precision and recall in a highly skewed dataset is quite notable. Thus, LinkDTI is able to integrate heterogeneous biological data and effectively model complex interactions in computational drug repurposing and drug discovery. Since scalability is a common setback in knowledge graph networks, expanding the scalability of the model can be a promising topic for future research.

---

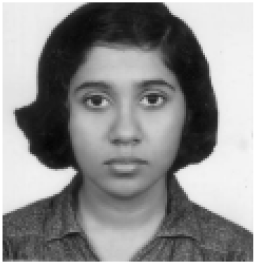

**MADHURIMA MONDAL** is a Ph.D. student in the Genomic Signal Processing Lab at the Department of Electrical and Computer Engineering at Texas A&M University, College Station, USA. She did her bachelor’s and master’s in Electronics and Electrical Communication Engineering from the Indian Institute of Technology, Kharagpur in India. Her research focuses on Deep learning applications in bioinformatics, with a particular interest in drug repurposing and network modeling for cancer

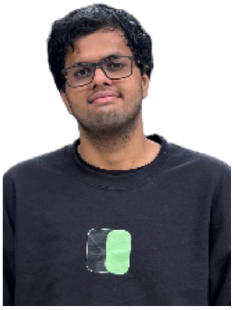

**SUBRAHMANYAM ARUNACHALAM** is an AI Practitioner and Researcher with interests in Reinforcement Learning, Computer Vision and Generative AI. He received his M.S degree in Computer Engineering from Texas A&M University, College Station, USA and B.Tech degree in Instrumentation and Control Engineering from National Institute of Technology, Tiruchirapalli, India. His motivations are applying Machine Learning techniques to realworld problems.

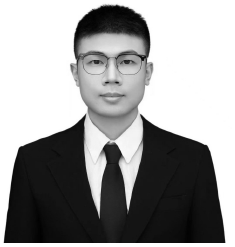

**SHILI WU** is a Ph.D. student in the Genomic Signal Processing Lab at the Department of Electrical and Computer Engineering at Texas A&M University, College Station, USA. He received his B.S. degree from Case Western Reserved University and M.S. degree from Columbia University. His research interests includes reinforcement learning and control application. He is also interested in applying the machine learning techniques into biography. He has authored several papers in these areas and is an IEEE member.

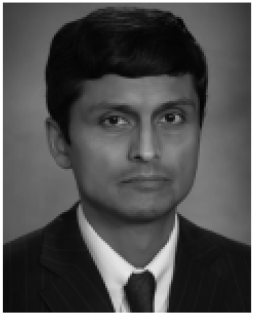

**ANIRUDDHA DATTA (FELLOW, IEEE)** received the BTech degree in electrical engineering from the Indian Institute of Technology, Kharagpur, India, in 1985, the MSEE degree from Southern Illinois University, Carbondale, Illinois, in 1987, and the MS (applied mathematics) and PhD degrees from the University of Southern California, Los Angeles, California, in 1991. In August 1991, he joined the Department of Electrical and Computer Engineering, Texas A&M University where he is currently the J. W. Runyon, Jr. ‘35 professor II and Co-Director of Graduate Programs. His research interests include adaptive control, robust control, PID control, and Genomic signal processing. He has authored or coauthored five books and more than 200 journal and conference papers on these topics. He has served as an associate editor of the IEEE Transactions on Automatic Control (2001-2003), the IEEE Transactions on Systems, Man and Cybernetics-Part B (2005-2006), the IEEE Transactions on Biomedical Engineering (2013-2015), the EURASIP Journal on Bioinformatics and Systems Biology (2007-2016), the IEEE Journal of Biomedical and Health Informatics (2014-2016), the IEEE/ACM Transactions on Computational Biology and Bioinformatics (2014-2017), the IEEE Access (2013-2022) and is currently serving as an Academic Editor for PLOS One.

